# How the layer-dependent ratio of excitatory to inhibitory cells shapes cortical coding in balanced networks

**DOI:** 10.1101/2024.11.28.625852

**Authors:** Arezoo Alizadeh, Bernhard Englitz, Fleur Zeldenrust

**Affiliations:** Donders Institute for Brain, Cognition, and Behaviour, Radboud University, Nijmegen, the Netherlands

## Abstract

The cerebral cortex exhibits a sophisticated neural architecture across its six layers. Recently, it was found that these layers exhibit different ratios of excitatory to inhibitory (EI) neurons, ranging from 4 to 9. This ratio is a key factor for achieving the often reported balance of excitation and inhibition, a hallmark of cortical computation.

Although previous studies have explored various biologically realistic aspects of classical balanced networks, such as the effects of more complex neuronal properties including spike frequency adaptation using adaptive exponential integrate-and-fire (AdEx) neuron models (1), the temporal evolution of randomly connected balanced networks through second-order mean-field approximations (2), predictions of nonlinear dynamics across diverse neuron models ranging from integrate-and-fire to Hodgkin–Huxley formulations, thus bridging population dynamics to neuronal electrophysiology (3), the impact of spatial heterogeneity in balanced networks (4–6), and variations in synaptic timescales (7), the specific effects of differences in excitatory-inhibitory (EI) ratios on layer-specific dynamics and computational properties have, to the best of our knowledge, not yet been addressed. We investigate this question using a sparsely connected network model of excitatory and inhibitory neurons. To keep the network in a physiological range of firing rates, we varied the inhibitory firing threshold or the synaptic strength between excitatory and inhibitory neurons. We find that decreasing the EI ratio allows the network to explore a higher-dimensional space and enhance its capacity to represent complex input. By comparing the empirical EI ratios of layer 2/3 and layer 4 in the rodent barrel cortex, we predict that layer 2/3 has a higher dimensionality and coding capacity than layer 4. Furthermore, our analysis of primary visual cortex data from the Allen Brain Institute corroborates these modeling results, also demonstrating increased dimensionality and coding capabilities of layer 2/3.

**Author summary:** Experimental studies indicate that the ratio of excitatory to inhibitory neurons varies across different cortical layers. In this study, we investigate how these varying excitatory-to-inhibitory (EI) ratios affect the layer-specific dynamics and computational capacity of cortical networks. We modeled a randomly connected network of spiking neurons, incorporating different EI ratios based on experimental observations. Our findings reveal that as the influence of inhibition increases, corresponding to lower EI ratios, the network explores a higher dimensionality in its activity, thereby enhancing its capacity to encode high-dimensional inputs. These results align with our analysis of experimental data recorded from layers 2/3 and layer 4 of the rodent primary visual cortex. Specifically, our findings support the hypothesis that layer 2/3, which has a lower EI ratio compared to layer 4, possesses a greater computational capacity.

## Introduction

The activity recorded from neurons in the cortex of behaving animals shows temporally irregular spike patterns from a single neuron perspective, asynchronous activity from a population perspective, and a large trial-to-trial variability (8,9). Theoretical studies proposed that this asynchronous and irregular activity can be explained by a tight balance between excitation and inhibition in randomly and sparsely connected networks (10–14), which has become one of the corner-stone theoretical frameworks for understanding cortical dynamics. If inhibition in the balanced network is strong enough to counterbalance excitation, the total mean of the average input to each neuron is just below the threshold and neural action potentials are generated by fluctuations around this mean. As a result, the overall output patterns are irregular. Irregular outputs lead to irregular inputs, and the recurrent nature of the connections in the network maintains the neuronal activity in an asynchronous state.

The existence of the balance between excitation and inhibition described in the previous paragraph has been validated by several experimental measurements *in vivo*. In the context of cortical upstate oscillations, Compte et al. (2009) and Wehr et al. (2003) (15,16) demonstrated that the number of detected inhibitory events tracked the number of excitatory ones. In turn, Dehghani et al. (2016) (17) showed that during ongoing cortical activity and sleep, there is a balance characterized by a qualitative match between the population spike rate histograms of excitatory and inhibitory neurons. By examining intracellular synaptic potentials of multiple units and local field potentials during the up state of slow oscillations in the prefrontal cortex *in vivo* (18), it was found that excitatory and inhibitory conductances are proportional and well-balanced during periods of spontaneous network activity in the intact neocortex. On a single neuron level, Cafaro et al. (2010) and Graupner et al. (2013) (19,20) showed that excitatory and inhibitory inputs are correlated on a millisecond time scale.

A key property for achieving a balanced state is the network composition, i.e. the ratio and relative synaptic output strength of the excitatory and inhibitory neurons, both in biological and simulated neural networks. Previous theoretical studies and simulations mostly assumed a ratio of excitatory to inhibitory (EI) numbers of neurons, N_exc_/N_inh_ = 4 (21–23), based on the seminal work of Abeles et al. (1991) (24), which we will refer to here as the ‘classical’ balanced states. However, recent studies have shown that the EI ratio across different cortical layers can differ substantially from this ratio, in fact ranging from 4-9 (25–34). Huang et al. (2022) (26) employed *in vitro* fluorescence imaging to quantify cell bodies across six cortical layers of the barrel column through immunochemical labeling and confocal microscopy. The findings revealed substantial differences in the densities of different cell populations across these layers. Specifically, an EI ratio of 5.25 in layer 2/3 and 7.34 in layer 4 was found (see Fig. 1), supporting the conclusion that EI ratios differ between layers and areas (35–37).

**Fig. 1.**
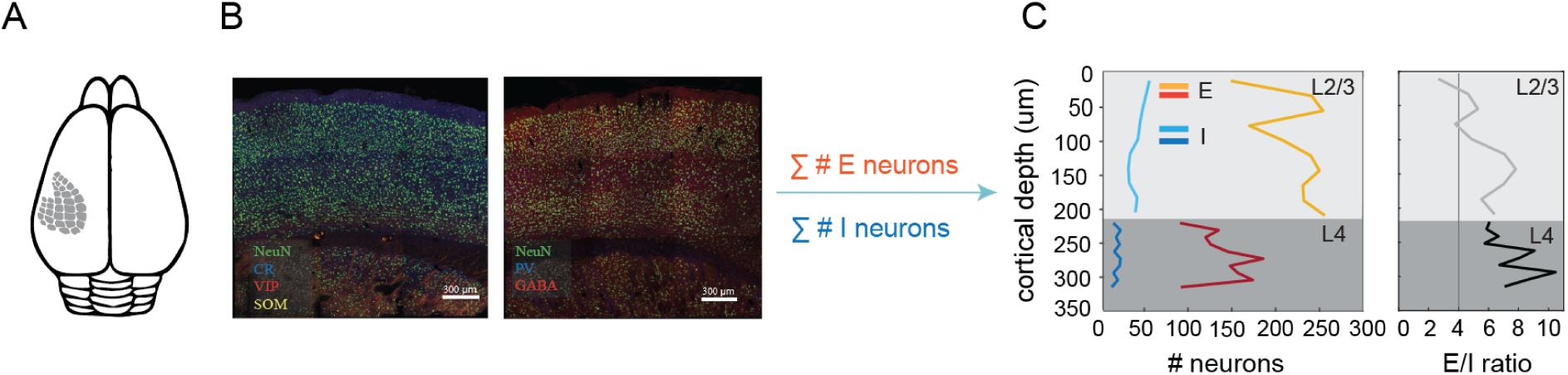
The cellular density distribution and excitatory-inhibitory balance vary across layers in the mouse barrel cortex. A) Schematic representation of the recorded area. B) Immunohistochemical staining using markers for inhibitory neurons (GABAergic, somatostatin-positive) and excitatory neurons (calretinin-, VIP-positive). C) Quantitative analysis of neuronal density in the barrel column. Cell bodies were quantified across layer 4 with cortical depth 440-629 µm and layer 2/3 with cortical depth 22-418 µm using immunochemical labeling and confocal microscopy. Red and yellow colors represent excitatory neuron density and light and dark blue represent inhibitory neuron density across layers 4 and layer 2/3 respectively. The final column demonstrates the relative ratio of excitatory to inhibitory neurons in each layer. Key findings include a variation in the number of excitatory and inhibitory neurons across cortical depth (418-629 µm) and a changing ratio of excitatory to inhibitory neurons as a function of cortical depth. A solid vertical line demonstrates the EI ratio in the classical balanced state. Figure adapted from Huang et al. (2022) (26) with permission.

Even though different EI ratios have been reported recently (26), so far, no study has addressed how these differences in layer-specific EI ratio will affect layer-specific dynamics and computational properties. Here, we investigate this question by varying the EI ratio in a randomly-connected network of excitatory and inhibitory spiking neurons (21). We consider different EI ratios in the balanced network by varying the number of inhibitory neurons. To keep the network in a physiological operating range in terms of firing rate, we either varied the firing threshold of the inhibitory neurons (which has been shown to differ between cortical excitatory and inhibitory neurons, see Fig. S1, based on data from (38)) or the synaptic strength from inhibitory to excitatory neurons.

We investigate the relation between the EI ratio, the dimensionality of the network response and the coding capacity of the network, by measuring the participation ratio and computational capacity of the network with different EI ratios as well as decoding temporally varying inputs. We find that as the influence of inhibition in the network increases, by either decreasing the EI ratio, increasing the strength of the inhibitory on excitatory activity, or decreasing the threshold of inhibitory neurons, the network explores a higher dimensionality in its activity, which increases the network’s capacity to represent high dimensional inputs. By mimicking the measured properties of layer 2/3 and layer 4 of the rodent barrel cortical networks (26), we conclude that the coding capacity of layer 2/3 is likely higher than that of layer 4, and corroborate this by performing a comparable analysis on experimental data from the Allen Brain Institute (39).

## Results

To understand how the EI ratio influences the cortical layers’ dynamics and computations, we built on a well-established network model ((21), see Methods) containing excitatory and inhibitory leaky integrate-and-fire (LIF) neurons. This model’s simplicity allows for an in-depth analysis with minimal complexity. We fixed the number of excitatory neurons (N_exc_) at 10,000 while systematically reducing the number of inhibitory neurons (N_inh_) from 2,500 to 1,000 with steps of 150. This manipulation creates different EI ratios within the network, ranging from 4 to 10, sampling the experimentally observed variation in EI ratios across different cortical depths in the barrel cortex (26). To maintain biologically relevant firing rates despite the increasing EI ratio, we adjusted the inhibitory influence on the network in two ways: either by strengthening the inhibitory connections to excitatory neurons (J_ei_) or by lowering the threshold at which inhibitory neurons fire (θ_inh_), as inhibitory neurons inherently have lower thresholds compared to excitatory neurons (21, Fig. S1). In essence, we fine-tuned three key network parameters to explore how the EI balance affects cortical layers’ dynamics, as illustrated in Fig. 2B.

**Fig. 2:**
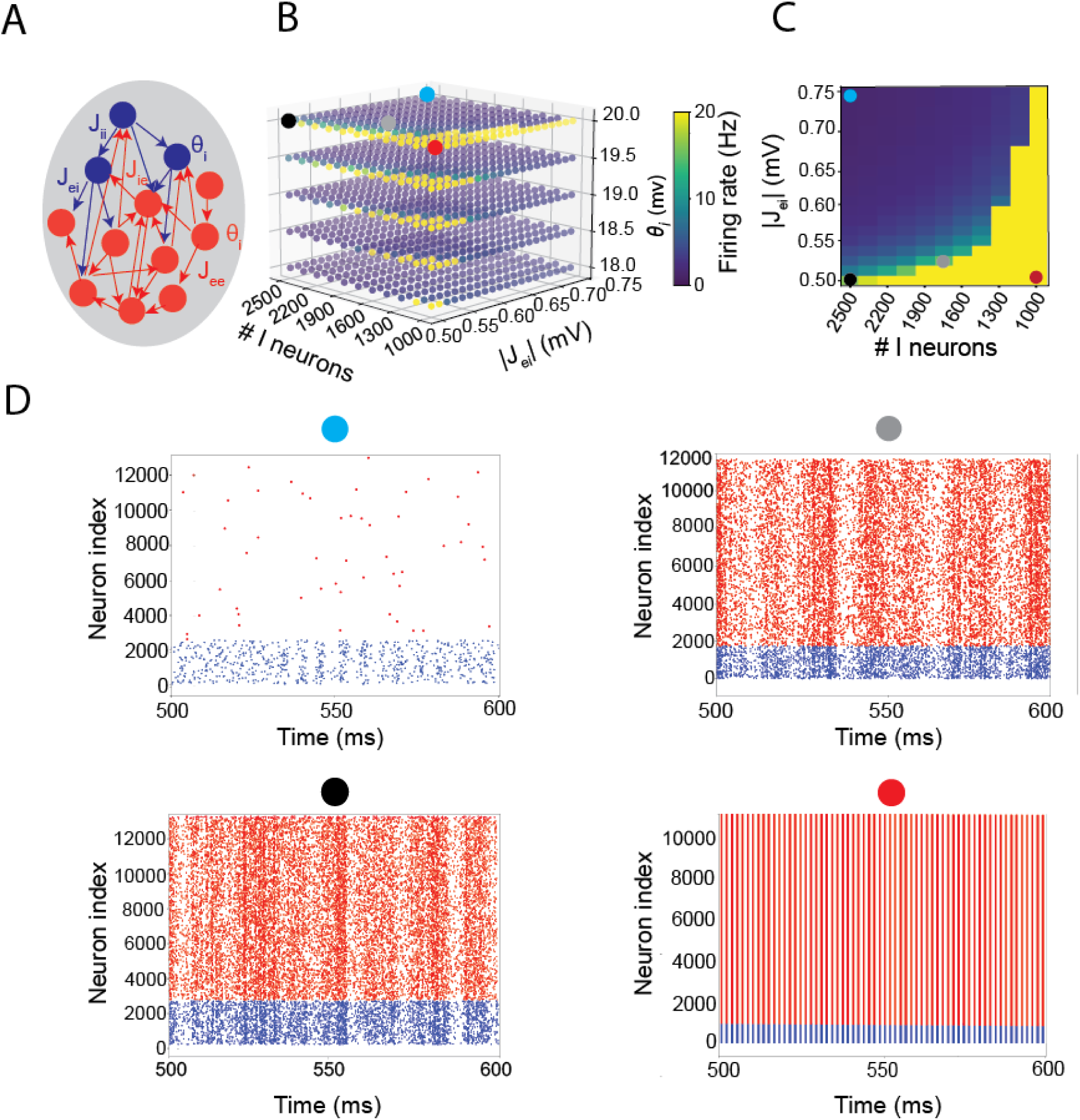
Spiking neural network diagram and activity. **A)** The network model consists of excitatory and inhibitory neurons that are randomly connected. We explored the effects on network dynamics and computation of the following variables: the number of inhibitory neurons N_inh_ (between 2500 and 1000), and consequently the EI ratio (between 4 and 10), the synaptic coupling strength J_ei_ (between −0.75 mV and −0.5 mV), the inhibitory firing threshold θ_inh_ (between 18 mV and 20 mV). Further, we used a fixed number of excitatory neurons (N_exc_ = 10,000), external input (μ_0_ = 24 mV) and probability of connection (10%). **B)** Averaged firing rate of the neurons in the network across EI ratios, J_ei_, and θ_inh_. Four circles (black, gray, red, and blue) demonstrate the network dynamics in four different states: black and grey: balanced excitation-inhibition dynamics (black circle: EI ratio = 4, J_ei_ = −0.5 mV, θ_inh_ = 20 mV (21), grey circle: EI ratio = 5.71, J_ei_ = −0.525 mV, θ_inh_ = 20 mV), red: excitation-dominated dynamics (EI ratio = 10, J_ei_= −0.5 mV, θ_inh_ = 20 mV), blue circle: inhibition-dominated dynamics (EI ratio = 4, J_ei_ = −0.75 mV, θ_inh_ = 20 mV) **C)** Averaged firing rate of the neurons in the network across EI ratios, J_ei_, and θ_inh_ = 20 mV with the same four dots demonstrating four states mentioned in B. **D)** Corresponding raster plots of the four states in B. Red dots indicate excitatory spikes and blue dots inhibitory spikes.

### Network activity with constant input (resting state)

First, we assess the network dynamics without external input. We refer to this as the ‘resting state’ of the network, the state when there is no time-dependent input and the network is driven by a constant offset current (22). Increasing the EI ratio, which can be achieved by decreasing the number of inhibitory neurons N_inh_ in the network (Fig. 2B, red circle), results in the excitatory neurons dominating the activity, so the network settles into a strongly synchronized state with regularly firing neurons behaving as oscillators (Fig. 2D, the raster plot with a red circle). As the inhibitory drive increases, either by increasing J_ei_ (Fig. 2B, blue circle), N_inh_ or by decreasing θ_inh_, the inhibition starts to dominate the excitation, resulting in strongly dampened excitatory firing at low rates (Fig. 2D, the raster plot with blue circle). In this state, the global network firing rate becomes stationary and remains very low. The classical asynchronous state occurs for parameters N_inh_ = 2500, J_ei_ = −0.5 mV and θ_inh_ = 20 mV (Fig. 2B and 2D raster plot, black circle) (21). To keep the network in this balanced state while reducing the number of inhibitory neurons, the inhibitory drive and the network firing rate can be kept constant by compensating the reduction of inhibitory neurons by increasing J_ei_ (Fig. 2B, grey circle) or θ_inh_.

### Effects of changes in EI ratio on network dynamics

Different EI ratios in the resting state imply vastly different computational properties. To further illustrate these properties, we examined the dynamics of the network in response to two different time-dependent inputs (Fig. 3A) for different values of the EI ratio, J_ei_, and θ_inh_. To assess the network dynamics in response to these two inputs, we assessed 3 properties of the network activity:

1. We measured the regularity of the spiking patterns by the coefficient of variation of the interspike-interval distribution (CV_ISI_) (Fig. 3B)
2. We measured the synchrony of the network activity by averaging the cross-correlations between neuron pairs’ spike trains over neuron pairs, normalizing the cross-correlation values between 0 and 1 (Fig. 3C).
3. We assessed the dimensionality of the network dynamics by computing the participation ratio (PR) (see Methods, (40)).

**Fig. 3:**
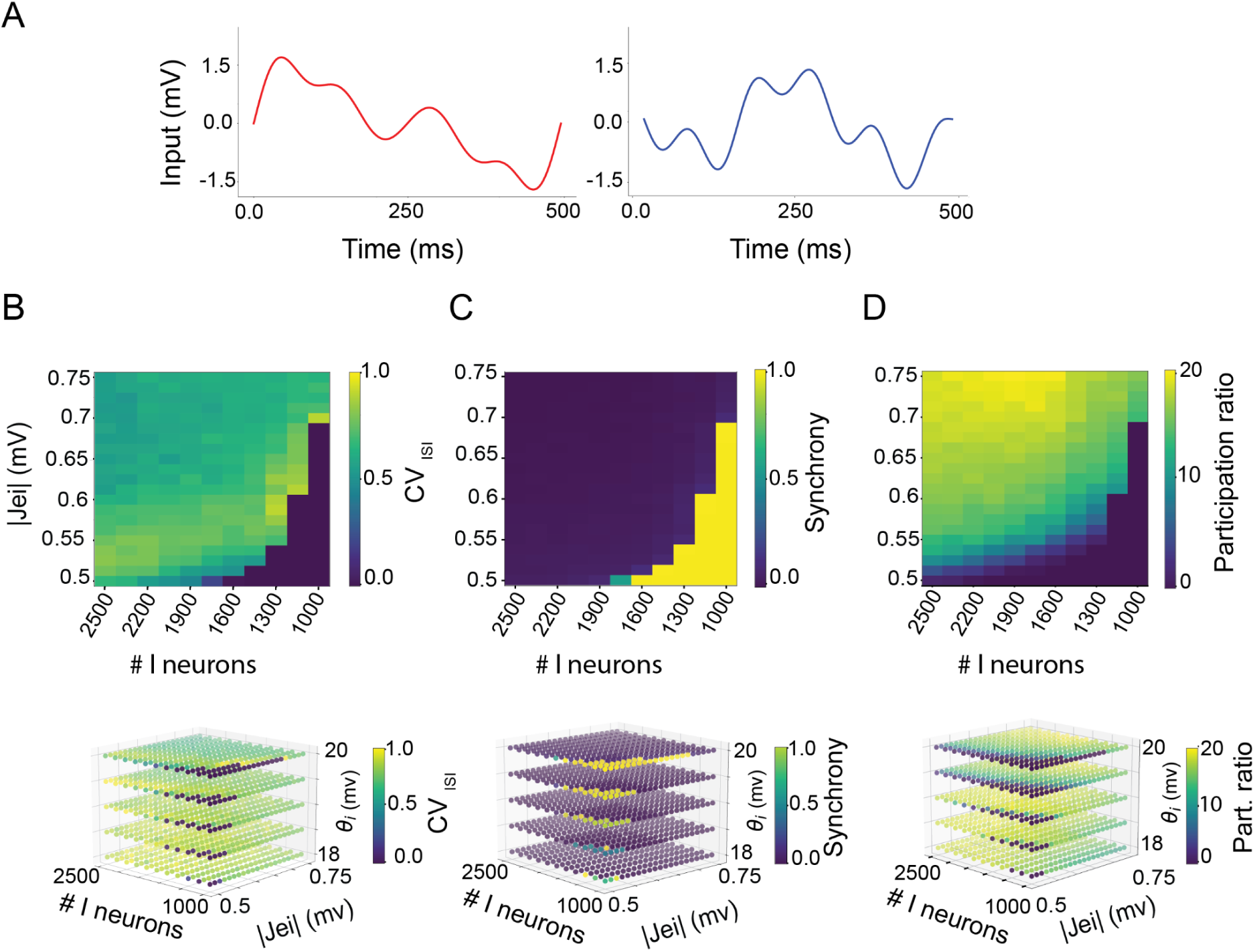
The network activity across varying EI ratios, J_ei_ and θ_inh_ explores a high-dimensional space when inhibition dominates. **A)** Two different temporally varying inputs, stimulus 1 and stimulus 2 were given to the network. **B)** top: Average coefficient of variation of inter-spike intervals (CV_ISI_) averaged over the neurons in the network for the following parameters: the number of inhibitory neurons N_inh_ between 2500 and 1000, and consequently the EI ratio between 4 and 10, J_ei_ between −0.75 mV and −0.5 mV, and θ_inh_ = 20 mV. Bottom: Same as top, but for θ_inh_ values between 18 mV and 20 mV. **C)** top: Normalized averaged synchrony between all neuron pairs in the network measured by cross-correlation for the same EI ratios, J_ei_, and θ_inh_ as in A. Bottom: Same as top, but for the θ_inh_ between 20 mV and 18 mV. **D)** top: Average participation ratio (PR) of the network for the same parameters: EI ratio, J_ei_, and θ_inh_ as in A. Bottom: Same as top, but for θ_inh_ between 18 mV and 20 mV.

All measurements were averaged over all neuron pairs, for all EI ratio values between 4 and 10 individually, all J_ei_ between −0.75 mV and −0.5 mV individually, and θ_exc_ of 20 mV (Fig. 3B-D, top Figs) and all θ_inh_ between 18 mV and 20 mV individually (Fig. 3B-D, bottom row). Using these three measurements, we find that the network’s dynamics depend on the inhibitory drive. In principle, the firing rate of every neuron is a potential degree of freedom in the dynamics of the network, and the response to an input can be represented as a trajectory in a high-dimensional space where every dimension represents the activity of one neuron (34, 35). In the synchronized regular regime, when excitation dominates the network (14, 36), the responses of all neurons are highly similar, the dynamics effectively explore only one dimension of this huge space. The dimensionality of the activity of the network generated by two inputs (Fig. 3D) indeed confirmed that the network’s dynamics could be reduced to a single dimension in the synchronized, regular regime.

When the EI ratio, θ_inh_ and J_ei_ are adjusted to strengthen the inhibitory drive to the point where it cancels out excitation, the CV_ISI_ abruptly rises to values greater than 0.5 (Fig. 3B) and the synchrony drops to values below 0.5 (Fig. 3C). This indicates highly variable inter-spike intervals and a low synchrony between neurons, coinciding with a sharp decrease in firing rates. In such cases, the network operates in the Asynchronous Irregular (AI) state (21) in which temporally varying inputs can interact with the internal state of the network to produce complex dynamics (7, 37). The dynamics of the network explore many more dimensions (Fig. 3D). Essentially, the balance between excitation and inhibition determines the network’s operational mode and how many dimensions its activity can explore. Next, we will assess how this influences the network activity for parameter values that are realistic for barrel cortex layers 4 and layer 2 /3.

### Network coding capacity for realistic cortical EI ratios

Given the critical role of the EI ratio on the excitation-inhibition balance, which shapes the network dynamics, we investigated how these dynamics vary across cortical layers. We measured the dimensionality of our neural network model by simulating a network that has properties mimicking those of layers 4 and 2/3 of the barrel cortex. According to our recent work (26) (Fig. 1), layer 4 has an EI ratio of 7.35 and layer 2/3 has an EI ratio of 5.25. The firing rate of neurons in layer 4 naturally depends on the type of input they receive. However, it has been estimated to be at least twice that of layer 2/3 (45). Finally, we know that the thresholds of inhibitory neurons are lower, and their firing rates higher (21). We investigated the dimensionality of the network activity for EI ratios similar to those found in the barrel cortex (EI ratio = 7.7 for layer 4, EI ratio = 5.3 for layer 2/3) (26) and maintained realistic firing rates by adjusting J_ei_ or θ_inh_ while preserving the firing rate differences between these two layers. Due to the significant increase in network inhibition caused by reducing θ_inh_, it was challenging to find consistent firing rates across all θ_inh_ values. To address this, we focused on θ_inh_ values of 20 mV, 19.5 mV, and 19 mV and selected firing rates of 8 Hz, 4 Hz, and 2 Hz for the analysis. For a fixed θ_inh_, we sought J_ei_ values that preserved constant firing rates across all EI ratios, effectively compensating for variations in inhibitory drive caused by changes in the EI ratio (Fig. S3). The highest consistent firing rate observed was 8 Hz for θ_inh_ values of 20 mV and 19.5 mV. To maintain the network’s firing rate at 8 Hz with an EI ratio of 7.35 (mimicking layer 4) and at 4 Hz with an EI ratio of 5.25 (mimicking layer 2/3), we adjusted the inhibitory synaptic strength (J_ei_) for different θ_inh_ values (Fig. 4, black dots). Similarly, to achieve firing rates of 4 Hz with EI ratio = 7.35 (mimicking layer 4) and at 2 Hz with an EI ratio of 5.25 (mimicking layer 2/3), different J_ei_ values were required for each EI ratio and θ_inh_ (Fig. 4, red dots).

**Fig. 4:**
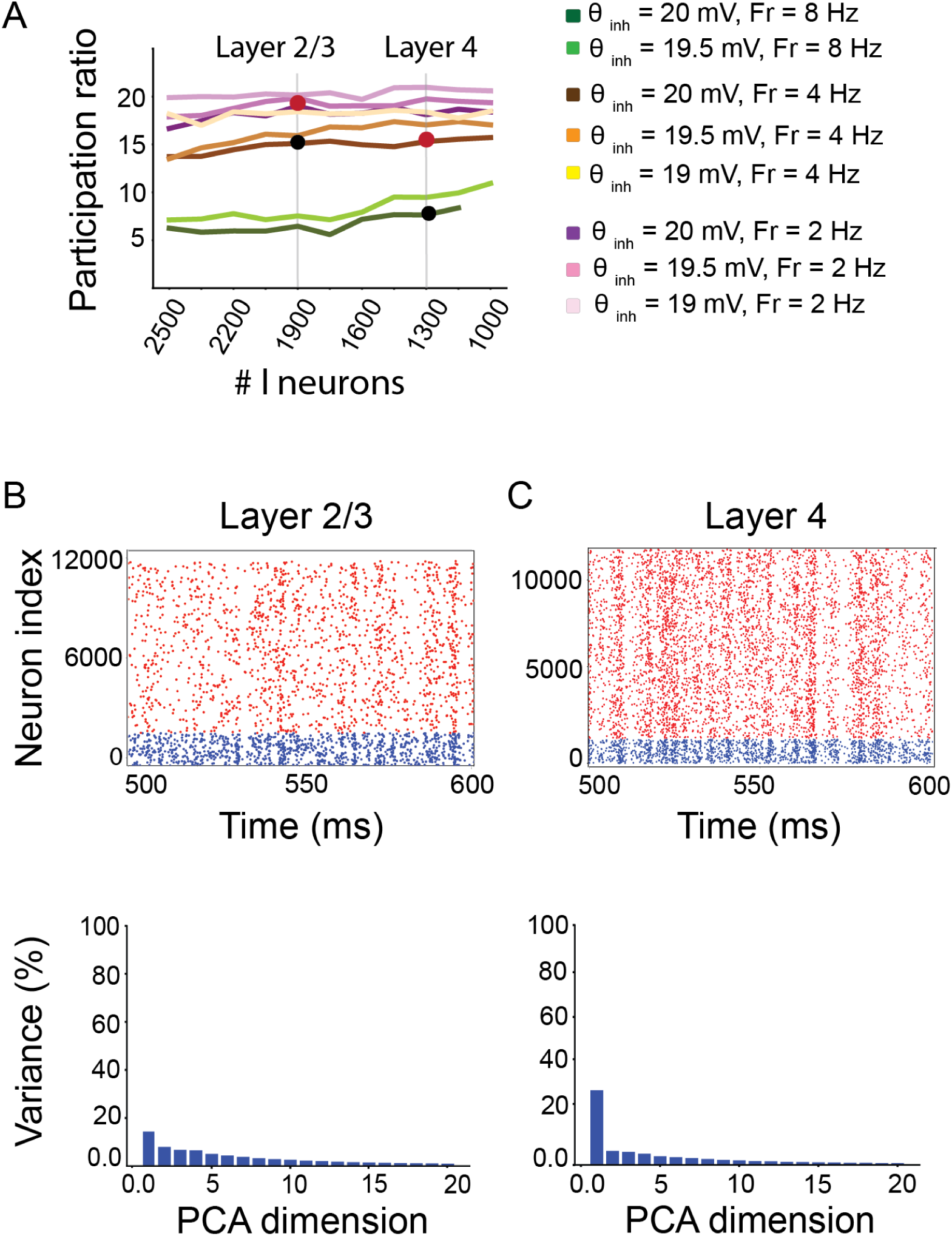
Networks mimicking the properties of cortical layers 2/3 (B) and layer 4 (C) respectively, show a difference in response dimensionality. **A)** The participation ratio of the network at averaged neuron firing rates of 8 Hz, 4 Hz, and 2 Hz as a function of the EI ratio (the number of inhibitory neurons), and θ_inh_. For all θ_inh_ and EI ratios, J_ei_ were adjusted to maintain the average firing rate of the network at these three values. Vertical lines at EI ratios of 5.3 (N_inh_= 1900) and 7.7 (N_inh_ = 1300) represent layers 2/3 and layer 4 of the barrel cortex with different θ_inh_, firing rates, and J_ei_. We compared layer 2/3 with layer 4 at two different firing rates: Black dots: Layer 2/3 (EI ratio = 5.3, θ_inh_ = 20 mV, J_ei_ = −0.575, firing rate = 4 Hz) and layer 4 (EI ratio = 7.7, θ_inh_ = 20 mV, J_ei_ =-0.575 mV, firing rate = 8 Hz). Red dots: Layer 2/3 (EI ratio = 5.3, θ_inh_ = 20 mV, J_ei_ = −0.7125, firing rate = 2 Hz) and layer 4 (EI ratio = 7.7, θ_inh_ = 20 mV, J_ei_ = −0.675, firing rate = 4 Hz). **B)** Top: A raster plot of layer 2/3 neurons, matching the left black dot in panel A with EI ratio = 5.3, PR = 15.1, firing rate = 4 Hz, and θ_inh_ = 20 mV. Blue and red dots represent spike times of excitatory and inhibitory neurons, respectively. Bottom: A principal component analysis (PCA) quantifies the dimensionality of the neurons in this model of layer 2/3. **C)** Same as B, but for a model mimicking layer 4 with EI ratio = 7.7, PR = 7.6, firing rate = 8 Hz, and θ_inh_ = 20 mV, right black dot in panel A.

As the average neuron’s firing rate decreases from 8 Hz to 2 Hz, due to increased inhibition, the network activity’s dimensionality increases (Fig. 4A; 8 Hz: green lines, 2 Hz: purple lines). However, for a given firing rate, the dimensionality remains almost the same across the different EI ratios and fixed θ_inh_ (Fig. 4A). The increase in the network dimensionality is larger when network firing rate decreases from 8 Hz to 4 Hz for all EI ratios and θ_inh_= 20mV and 19.5 mV (Fig. 4A, compare green and brown lines) than when it decreases from 4 Hz to 2 Hz (Fig. 4A, compare brown and purple lines) for all EI ratios and θ_inh_ = 19.5 mV and 19 mV, indicating that the network has a limited capacity of increasing its activity’s dimensionality and will eventually become saturated.

Interestingly, under constrained conditions where both inhibitory population size (Fig. 4A N_inh_ = 1900) and mean firing rate (Fig. 4A 4 Hz; brown lines) were fixed, a reduction in θ_inh_ (dark to light gradients) necessitated a decrease in J_ei_ to maintain firing rate fixed. Under these iso-rate conditions, we observed an increase in network dimensionality. This result suggests that, even with a fixed firing rate, variations in inhibitory parameters can shape the network’s dynamics.

When comparing different putative ‘layers’ (i.e. using matched EI ratios and firing rates), we found that a network configuration to resemble layer 4 (EI ratio 7.7, firing rate = 8 Hz, θ_inh_ = 20 mV, J_ei_ = −0.575 mV) had a dimensionality of 7.6 (black dot in Fig. 4A), while a layer 2/3 configuration (EI ratio = 5.3, firing rate = 4 Hz, θ_inh_ = 20 mV, J_ei_ = −0.575) exhibited a dimensionality of 15.1 (black dot in Fig. 4A). We repeated the experiment with lower firing rates (layer 4: EI ratio = 7.7, firing rate = 4 Hz, θ_inh_ = 20 mV, J_ei_ = −0.675 mV; layer 2/3: EI ratio 5.3, firing rate = 2 Hz, θ_inh_ = 20 mV, J_ei_ = −0.7125), represented by red dots in Fig. 4A. Again, the layer 4 configuration showed lower dimensionality, (PR = 15.3), than the layer 2/3 configuration (PR = 19).

In addition to dimensionality, we assessed the synchrony and CV_ISI_ of the spike trains of the network models For the higher firing rate condition, layer 4 (EI ratio = 7.35, firing rate = 8 Hz) showed a CV_ISI_ of 0.69 and synchrony of 0.09, while layer 2/3 (EI ratio = 5.25, firing rate = 4 Hz) had a CV_ISI_ of 0.71 and synchrony of 0.036. At lower firing rates, layer 4 (EI ratio = 7.35, firing rate = 4 Hz) had a CV_ISI_ of 0.78 and synchrony of 0.05, whereas layer 2/3 (EI ratio = 5.25, firing rate = 2 Hz) had a CV_ISI_ of 0.76 and synchrony of 0.024.

In conclusion, we demonstrate that if we model networks to resemble what we know of different cortical layers (i.e. neurons in layer 2/3 exhibit sparser firing patterns and a lower EI ratio, inhibitory neurons have a lower threshold), the network activity of a network with layer 2/3 parameters occupies a higher-dimensional space than that of a network mimicking layer 4. This is evident in the broader distribution of the variance of the spiking activity across multiple principal components, as reflected in the PR. As the dimensionality of the network response has been correlated to a network’s capacity to classify different inputs (5,22), this indicates that neurons in layer 2/3 of the cortex are expected to be better capable of classifying different temporal inputs (Fig. 3A). We explore this relation between the dimensionality of the network response and the classification capacity further in the next section.

### Inhibition improves decoding performance of the neural network

A correlation between the complexity of the neural network activity and its computational power was recently proposed (5,22). Building on this, we examined how the EI ratio influences a network’s ability to process information. We trained an SVM decoder to distinguish between two different input patterns (Fig. 3A) based on the network’s neural activity. The SVM was initially trained on 80% of the data and then tested on the remaining 20%. We analyzed the SVM’s decoding performance across various EI ratios and inhibitory strengths of J_ei_. Given that inhibitory neurons typically have lower activation thresholds than excitatory ones (38), we set the θ_inh_ to 19.5 mV, slightly below the excitatory threshold θ_exc_ of 20 mV. Figs 5A and 5B demonstrate the dimensionality of the network response and the decoding performance across various EI ratios and J_ei_ values, with θ_inh_ = 19.5 mV.

**Fig. 5.**
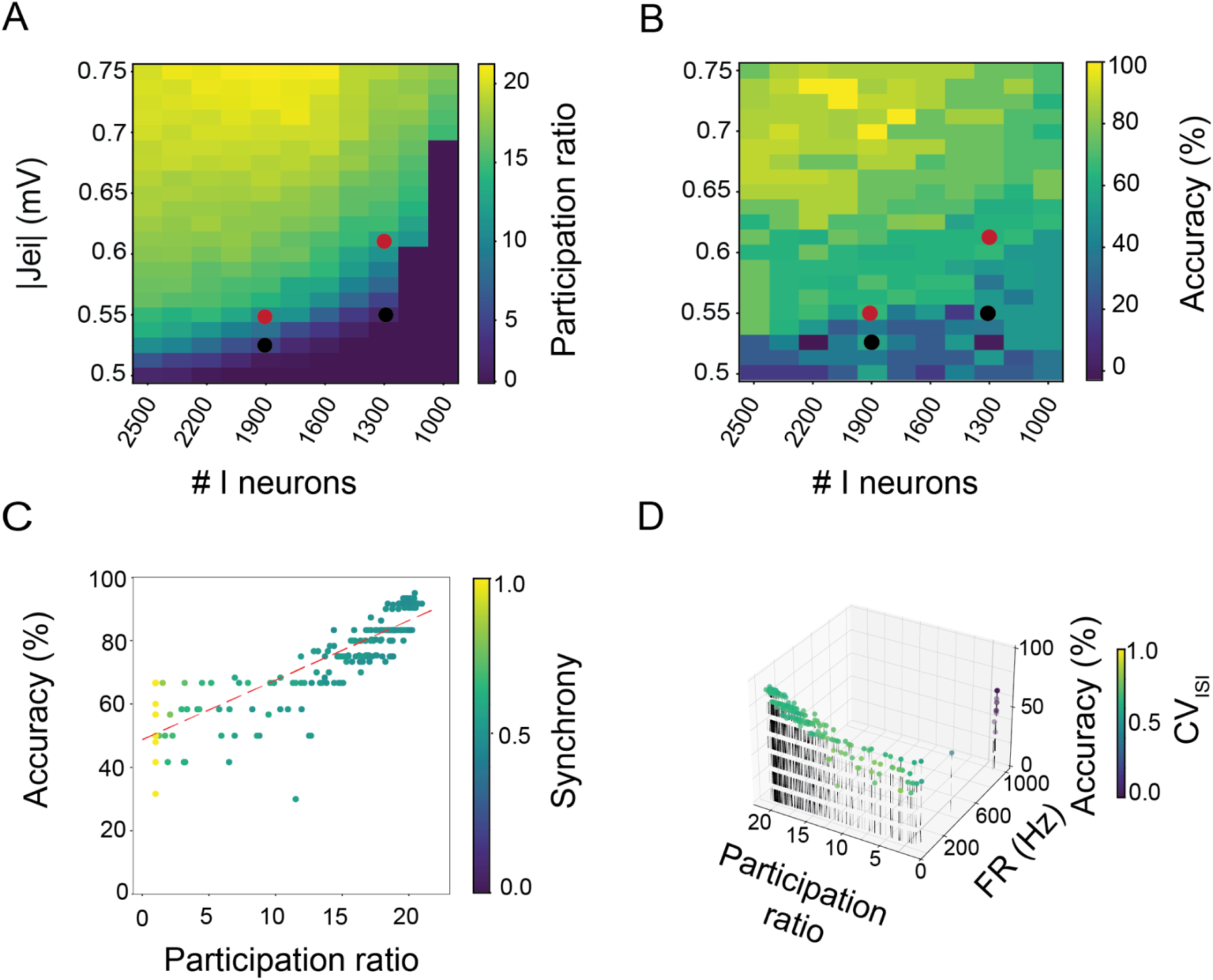
Correlation between response dimensionality and coding accuracy. A) Network dimensionality, measured as the the PR, as a function of the N_inh_ which determines EI ratio, J_ei_ at an inhibitory threshold θ_inh_ = 19.5 mV, of the activity of the network responding to two temporal inputs. The black and red dots represent the parameter values corresponding to layer 4 and layer 2/3 of the barrel cortex. Black dots correspond to higher firing rates (layer 4: 8 Hz, EI ratio of 7.7, J_ei_ = −0.55 mV; layer 2/3: 4 Hz, EI ratio of 5.3, J_ei_ = −0.525 mV), while red dots represent lower firing rates (layer 4: 4 Hz, EI ratio of 7.7, J_ei_ = −0.6125; layer 2/3: 2 Hz, EI ratio of 5.3, J_ei_ = −0.55) B) Network encoding capacity, evaluated using an SVM classifier to decode two temporal input stimuli from network activity, as a function of the EI ratio and J_ei_ at θ_inh_ = 19.5 mV. The black and red dots correspond to the same parameter values as in panel A. C) Synchrony of the neurons in the network in relation to the PR and network encoding capacity (Accuracy). The red line represents a linear fit between the accuracy and participation ratio (slope: 1.9, intercept: 48.74, which have a correlation of 0.85 and a p-value of p < 0.001). D) Regularity of network spike patterns (CV_ISI_) in relation to the PR, firing rate (FR) and network encoding capacity (Accuracy).

The dimensionality of the network response and the decoding accuracy show a strong correlation (r = 0.85, p < 0.001, Fig. 5C). Specifically, in conditions where excitatory neurons dominate, a substantial portion of the network enters a synchronous spiking state, which reduces the network dimensionality, severely impairing its ability to encode distinct temporal inputs. Under such conditions, the network’s performance in distinguishing between two external inputs is close to chance level (50%, Fig. 5B). In the asynchronous state, when inhibition dominates, either by increasing the number in inhibitory neurons N_inh_ or their synaptic strength J_ei_, the network activity explores additional dimensions and the decoding accuracy improves. The black and red dots in Fig. 5A and 5B represent the parameter values that we hypothesize correspond to layer 4 and layer 2/3 of the barrel cortex under different conditions (compared to Fig. 4). Black dots correspond to higher firing rates (layer 4: 8 Hz, EI ratio of 7.7, J_ei_ = −0.55 mV; layer 2/3: 4 Hz, EI ratio of 5.3, J_ei_ = −0.525 mV), while red dots represent lower firing rates (layer 4: 4 Hz, EI ratio of 7.7, J_ei_ = −0.6125; layer 2/3: 2 Hz, EI ratio of 5.3, J_ei_ = −0.55). In both conditions, layer 2/3 neurons explore a higher dimensionality and show a larger coding capacity than layer 4 neurons, p < 0.01, two-tailed t-test (layer 4: 8 Hz - dimensionality 9.5, encoding 67% ± 5.5; layer 2/3: 4 Hz - dimensionality 15.9, encoding 77% ± 9.7; layer 4: 4 Hz - dimensionality 17, encoding 77% ± 12.88; layer 2/3: 2 Hz - dimensionality 20.2, encoding 83.33% ± 9.8).

The analysis reveals that the network’s ability to encode information efficiently increases with the participation ratio PR (r = 0.85, p < 0.001, Fig. 5C) which coincides with reduced firing rates and a low regularity of spike patterns CV_ISI_. Figs. 5D illustrates the CV_ISI_ as a function of the PR, firing rate, and encoding capacity of the network. In this figure, each dot represents a specific value of the EI ratios and the J_ei_ when the θ_inh_ is set to 19.5 mV.

### Coding capacity in different layers of visual cortex

In order to investigate the impact of layer-specific EI ratios on the layer’s dynamics and computational properties, we simulated the spiking neural networks with sparse, random connectivity, and concluded that layer 2/3 has a higher dimensionality of the network response and a higher coding accuracy than layer 4. To validate these simulation results, we incorporated experimental data into our analysis. We analyzed a publicly available dataset from the Allen Brain Institute that included recordings from 60 head-fixed mice presented with a standardized set of visual stimuli (22, see Methods for details). Neuronal activity was captured simultaneously from multiple brain regions, including the primary visual cortex VISp. Focusing on VISp neurons, selected for their rich visual content and demonstrated effectiveness in neural decoding (39,46), we investigated the influence of EI ratio variations across cortical layers 2/3 and layer 4. To this end, we analyzed 250 ms responses to natural scene stimuli from the Brain Observatory 1.1 dataset (39).

We classified neurons in VISp layers 2/3 and layer 4 as excitatory or inhibitory based on AP waveform duration (Fig. S2). To maintain consistent EI ratios with the barrel cortex as used in the analysis above, we repeatedly selected 181 excitatory and 34 inhibitory neurons from layer 2/3 neurons, and 111 excitatory and 15 inhibitory neurons from layer 4 neurons to obtain EI ratios of 7.4 and 5.32 for layers 4 and layer 2/3, respectively (Methods: Analyzing NeuroPixels Dataset).

To estimate the computational capacity of VISp layers 2/3 and layer 4, we assessed the dimensionality and encoding capabilities of the selected populations. The dimensionality was quantified using the PR as described in Eq. 6, while the encoding capacity was determined through SVM decoding of natural scenes (Methods: Analyzing Neuropixels Dataset). We find that layer 2/3 neurons exhibited significantly higher PR values (5.23±0.24) compared to layer 4 (4±0.04), with p < 0.01 suggesting a higher dynamical dimensionality, in agreement with our model results. Consistent with this, SVM decoding accuracy for natural scenes was significantly higher in layer 2/3 (0.1±0.063) than in layer 4 (0.065±0.036), (p < 0.05, two-tailed t-test, N= 118) in both cases.

In addition to the dimensionality and encoding capacity, we also assessed the synchrony and CV_ISI_ of both layers. Layer 2/3 exhibited a significantly lower synchrony (0.05 ± 0.01) compared to layer 4 (0.08 ± 0.02; *p* < 0.05, two-tailed t-test). In contrast, the regularity of interspike intervals was similar between the layers, with CV_ISI_ values of 0.090 ± 0.02 for layer 2/3 and 0.088 ± 0.03 for layer 4.

These findings corroborate our modelling results that layer 2/3 possesses a superior stimulus separation capacity (in this case for natural scenes), likely due to its increased dynamical dimensionality.

## Discussion

In this study, we investigated how layer-specific EI ratios influence the dynamical behavior and computational capabilities of corresponding cortical layers, as observed in experimental studies (26). To this end, we developed a spiking neural network model composed of randomly and sparsely connected excitatory and inhibitory neurons, and adjusted the EI ratio to match physiologically observed values for layers 4 and 2/3 of the mouse barrel cortex. To maintain realistic firing rates across network configurations, we varied two additional parameters: (1) the firing threshold of inhibitory neurons, motivated by experimental findings showing intrinsic differences between excitatory and inhibitory neurons (see Figure S1 and reference (28)), and (2) the synaptic strength from inhibitory to excitatory neurons. The latter is supported by both theoretical studies (21,22), and experimental work emphasizing the importance of asymmetric synaptic strength in preserving stability and balance in cortical networks. For instance, Xue, et. al (2014) (47) demonstrated that synaptic strength is differentially regulated across cell types to maintain excitation-inhibition balance, while Isaacson and Scanziani (2011) (36) reported systematic differences in the timing and strength of excitation-to-inhibition versus inhibition-to-excitation connections.

We find that as inhibitory influence increases—driven by changes in threshold, relative synaptic strength, and number of neuron—the network explores a higher-dimensional space, enhancing its capacity to represent inputs. Importantly, this dimensionality expansion occurs independently of shifts in mean firing rates, suggesting that inhibition enhances representational capacity not through simple gain control, but by decorrelating population activity.

By comparing network models that take the experimentally found EI ratios, thresholds and firing rates of layer 2/3 and layer 4 in the mouse barrel cortex into account, we conclude that layer 2/3 likely has a higher dimensionality and coding capacity than layer 4. These results are mirrored by an analogous analysis on a large scale data set from the visual cortex by the Allen Brain Institute.

### Comparison with previous literature

Other recent studies have highlighted the link between network dimensionality and coding capacity: For example, Ostoijc (2014) (22) examined the dynamics and computational capabilities of an unstructured network of sparsely connected excitatory and inhibitory spiking neurons. They demonstrated that as the synaptic coupling strength increases, the network dynamic undergoes a transition from a classical asynchronous state to a heterogeneous asynchronous state. The former exhibits a low dimensionality, characterized by neurons firing at a constant rate, and the high redundancy of the activity that limits more complex transformations of inputs. In contrast, the latter displays higher dimensionality with neurons firing at diverse, time-varying rates, enabling more complex computations, such as temporal input categorization, at the cost of reduced signal propagation reliability. Gao et. al (2015, 2017) (40,48) subsequently revealed that the computational capacity of a neural network is not solely determined by the number of neurons but rather by the dimensionality of its activity. Neural responses often reside in a lower-dimensional subspace, and it is within this subspace that critical information processing occurs.

Recanatesi et. al (2022) (49) measured the dimensionality of neural population recordings across several brain regions, including sensory areas and decision-related areas, as well as during spontaneous state and an engaged state. They showed that brain regions closer to sensory input, such as the thalamus, have lower dimensionality than those further away, like the visual cortex. Additionally, brain areas involved in complex cognitive functions, like the frontal cortex, exhibit higher dimensionality during specific tasks compared to regions involved in simpler processes, like the midbrain. This implies that higher dimensionality might be linked to more complex cognitive functions and representations within the brain.

While the studies mentioned above have examined various factors influencing network dimensionality and consequently computational capacity, the specific role of layer-specific excitatory-inhibitory neuron ratios in determining network dimensionality and computational capacity remained largely unexplored. Here, we investigated the impact of the EI ratio on dimensionality and coding capacity, while maintaining a constant inhibitory drive. We hypothesized that a higher-dimensional network response would enhance the ability to differentiate between stimuli, whereas lower dimensionality would limit the encoding capacity. Indeed, we found a strong correlation between the dimensionality of the network response and the decoding accuracy (Fig. 5). In the balanced state, where excitatory and inhibitory neurons are in equilibrium, each neuron in the network exhibits asynchronous and irregular activity. This adds a new dimension to the network, increasing its overall dimensionality. As a result, the network can transform one-dimensional temporal inputs into higher-dimensional representations, creating a foundation for complex spatiotemporal computations. As a result, neurons within the network could easily classify the two temporal inputs (Fig. 4A) by performing linear discrimination. Strikingly, there appears to be a local maximum of decoding capacity around the EI ratio of about 5.3 (Fig. 5), the EI ratio that was previously found in layer 2/3 of the barrel cortex (26).

To explicitly assess the computational capacity of layer 4 and layer 2/3 of the cortex, we chose parameters of the network (EI ratio, firing threshold of inhibitory neurons, relative firing rates) to mimic supragranular and granular layers of the barrel cortex (26). We found the network model with realistic parameter values of layer 4 and layer 2/3 exhibited asynchronous irregular activity, but with different dynamics. Supragranular networks nonlinearly projected input activity into a higher-dimensional space (higher PR) than granular layers, enhancing their ability to differentiate stimuli. We corroborated our results against existing experimental recordings of neural activity (39), and found similar results. Therefore, we hypothesize that supragranular and granular layers of the cortex have different computational roles, where supragranular layers perform more complex or higher-order computations, whereas granular networks, with their lower dimensionality, may prioritize reliable information transmission over nonlinear processing, primarily transferring sensory information from the thalamus to supragranular layers.

Paradoxically, Dahmen et al. (2020) (50) reported a lower participation ratio in layer 2 compared to layer 4 within the mouse visual cortex of the Allen Brain Observatory. However, our findings diverged due to critical methodological differences. Specifically, we subsampled excitatory and inhibitory neurons from VISp layers 2/3 and 4 to align our excitatory/inhibitory (E/I) ratios with those of the barrel cortex (7.4 for L4 and 5.32 for L2/3). In contrast, Dahmen et al. (2020) (50) included the entire population of neurons from layer 2 and 4. This differential sampling approach likely accounts for the observed discrepancies in participation ratios between layers.

Based on the findings of Adesnik (2018) (35), it is reasonable to conclude that neurons in Layer 2/3 (L2/3) of the mouse‘s primary visual cortex are likely to exhibit more diverse or higher-dimensional activity than those in Layer 4 (L4), and this is at least partly due to stronger recurrent inhibition within L2/3. The study showed that during optogenetically induced gamma oscillations, the balance of excitation and inhibition (E/I ratio) in L2/3 was significantly more tilted toward inhibition (E/I ≈ 0.18) compared to L4 (E/I ≈ 0.27), indicating that L2/3 circuits are under tighter inhibitory control. This strong local inhibition suppresses redundant or correlated activity, enabling more selective and sparse firing patterns, which in turn can increase the dimensionality of population activity. Moreover, L2/3 plays a distinct integrative role in cortical processing: it receives inputs not only from L4 but also from other cortical areas and sends outputs mainly to L5, positioning it as a key hub for complex intracortical computation. This contrasts with L4, which acts primarily as a feedforward layer receiving thalamic sensory input and distributing it to other layers with more uniform excitatory output and less specialized activity. The study suggests that the strong inhibition observed in L2/3 contributes to an inhibition-stabilized regime that enables nonlinear gain control, contextual modulation, and flexible computation—features associated with high-dimensional, heterogeneous neural representations. Therefore, the combination of stronger inhibition, integrative connectivity, and computational function supports the idea that L2/3 neurons exhibit more diverse activity patterns than those in L4.

Across species from drosophila (51), rodents (28,29), cats (39,52), to non-human primates and humans (31), roughly 70–80% of cortical neurons are excitatory and 20–30% inhibitory, with the hippocampus and cerebellum showing ∼85–90% excitatory neurons (28,53,54). This conserved EI ratio supports memory, multi-task learning, and flexible information processing (34,55,56).

In the barrel cortex, the layer-specific distributions reflect this pattern while highlighting functional specialization: Layer 1 is almost entirely inhibitory (∼90–95%), with very few excitatory cells (∼5–10%), functioning primarily as a modulatory layer (52,57). In layers 2/3, 4, 5, and 6, excitatory neurons dominate numerically (∼75–85%), while inhibitory neurons make up ∼15–25% and include PV+, SOM+, and VIP+ interneurons that regulate local circuits and pyramidal outputs (33,57–59). These excitatory and inhibitory neurons form interlaminar circuits that balance activity, stabilize network dynamics, and allow flexible signal propagation across the cortical column (33,57–59).

### Limitations and future directions

Our neural network model, while based on simplified features and minimal biophysical constraints, benefits from a small number of free parameters, enabling a detailed understanding of its dynamics. However, several factors could influence our conclusions about the network’s dynamic behavior and computational capacity, which should be explored further. Firstly, cortical networks often exhibit clustered connectivity patterns as a skeleton of stronger connections embedded in a sea of weaker ones (60). This structure likely plays a crucial role in network dynamics. While a fully random population structure was sufficient for implementing a range of tasks, specific tasks appeared to require a non-random structure that could be described by a small number of statistically-defined sub-populations. Dubreuil’s analysis (61) revealed that such a population structure enabled flexible computations and shaped the dynamical landscape of collective dynamics. This finding suggests that, in order to study network computations effectively, there is likely value in considering localized connectivity rather than random structures, as it allows the network to achieve both flexibility and precision.

We employed standard leaky integrate-and-fire (LIF) neuron models, which assume a linear relationship between input current and neuron voltage (before spike threshold). However, in more realistic models, the voltage response to input is nonlinear, and the neuron exhibits a refractory period during which it is less susceptible to firing. Additionally, real neurons often exhibit spike-frequency adaptation (38,62), where the firing rate decreases over time in response to sustained input. Incorporating these non-linear factors can influence the dynamics of neural networks (38,62), which likely enriches the network activity.

A third simplification that might influence the results is the heterogeneity of neural properties. The brain is a complex network of interconnected neurons which have diverse physiological and spiking properties. This neural heterogeneity significantly influences brain dynamics and contributes to neural computations. For example, Zeldenrust et al., (2021) demonstrated that networks with diverse neuronal filters can represent input stimuli more efficiently. Perez-Nieves et al., (2021) found that individual membrane and synaptic time constants enhance task performance. Gast’s theoretical framework (5) discovered that the spike threshold heterogeneity of inhibitory neurons allows for a better gating of neural signals, maintaining balance, and preventing excessive activity. In excitatory neurons, spike threshold heterogeneity increases the dimensionality of neural dynamics, improving the network’s capacity for decoding tasks. In contrast, our spiking neural network models use uniform neural parameters across stimuli, resulting in a homogeneous network.

Fourth, we measured the dimensionality of the network response using the participation ratio PR (see Methods), a PCA-based, linear dimensionality estimation algorithm. A recent study (65) has evaluated various dimensionality estimation algorithms on synthetic datasets with known intrinsic dimensionality. While no algorithm is universally accurate, these assessments provide insights into when estimates are likely to be valid and when they are not. Linear methods, such as PCA, are generally less accurate than nonlinear methods when the mapping between the low-dimensional latent space and the high-dimensional space of neural recordings is nonlinear. However, PCA remains a popular choice due to its computational efficiency and ease of interpretation.

Fifth, the relation between network dimensionality and task performance likely depends on task dimensionality (33, 41). The task we used here, the discrimination of two stimuli, is relatively simple and low-dimensional. Different tasks should be explored to see how the relation between network dimensionality and task performance depend on various task parameters.

Finally, while both the barrel cortex and VISp follow general cortical principles, their wiring and organization differ substantially that might impact our comparison. The barrel cortex processes tactile input from whiskers via the VPM thalamus, forming distinct barrel structures in layer 4. In contrast, VISp processes visual input from the retina via the LGN, with a more uniform layer 4 lacking whisker-specific compartments (66). Although both regions exhibit columnar organization, the anatomical basis differs, as do the types and distributions of interneurons and horizontal connections tailored to their respective sensory roles (66). Plasticity mechanisms also vary accordingly (66–68). Due to the lack of suitable barrel cortex data, we used VISp data and adjusted the EI ratio to match our model. The similarity in dynamics suggests EI balance may be a fundamental circuit constraint, but differences in structure and function limit direct comparisons. Future work should aim to validate the model with actual barrel cortex data. Moreover, we recognize that incorporating more detailed features, such as inhibitory cell subclasses (SOM, VIP, PV), synapse-specific differences, realistic population sizes, and inter-layer connectivity, are essential for a more comprehensive understanding. We consider this work a foundation for subsequent investigations that will build upon these findings.

### Conclusions

In conclusion, we show that adapting balanced networks to three measured cortical network properties, the difference in EI ratio between cortical layers, the spiking thresholds of inhibitory neurons and the strength of the inhibitory connections to excitatory neurons, together result in different computational properties of supragranular and granular layer network models. A further exploration of the effects of biologically measured properties of neural networks, such as neural non-linearities, heterogeneity and the clustered structure of connectivity, should reveal the computational specialisation of different networks in the brain.

## Methods

### Network Simulations

#### Network of Spiking Neurons

The network model we used is identical to the classical network studied in (21). It consists of leaky integrate-and-fire neurons, with inhibitory and excitatory neurons simulated using the Brian2 spiking neural network simulator (69), randomly connected with a 10% robability of connection.

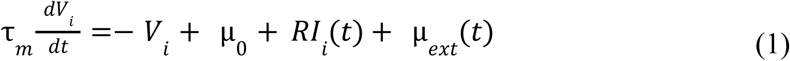

where *τ_m_* = 20 ms is the membrane time constant, µ_0_ = 24 mV is a constant offset current, *RI_i_* is the total synaptic input within the network and µ*_ext_*(*t*) is a time-dependent external current. When the membrane potential reaches the firing threshold *V*_th_ = 20 mV, an action potential is emitted and the membrane potential is reset to the resting potential V_r_ = 10 mV. The neurons’ dynamics resume after a refractory period of τ*_r_* = 0.5 ms.

The total synaptic input to the *i*_th_ neuron is:

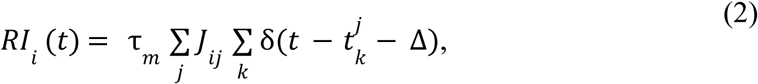

where *J_ij_* is the amplitude of the postsynaptic potential evoked in neuron *i* by an action potential occurring in neuron *j*, Δ = 0.55 ms is the synaptic delay and δ is a delta function representing a spike of a neuron *i* at time 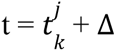.

In the ‘resting state’, each neuron receives a constant external input µ_0_ which effectively sets the resting membrane potential. There is no external noise, so the network is fully deterministic. When stimulated, a time-varying input, µ_ext_(t), is applied to 2,000 randomly selected excitatory neurons. This input consists of one of two stimuli, each a composite of superimposed sinusoidal waveforms. Each stimulus is presented for 500 ms in 30 trials, with each trial initiated by a 5 ms white noise perturbation to create diverse starting points.

The network consists of a fixed number of excitatory neurons, N_ext_ = 10000, and a variable number of inhibitory neurons to explore different EI ratios. The number of inhibitory neurons ranges from 2,500 (21) to 1,000 in steps of 150. Consequently, the EI ratio in the network varies between 4 and 10.

By maintaining the number of excitatory neurons constant and decreasing the number of inhibitory neurons in the network, the network’s firing rate experiences a significant increase. To ensure that the network remains within a physiological operating range in terms of firing rate, we decreased either the inhibitory firing threshold (θ_inh_) from 20 mV to 18 mV in steps of 0.5 mV, or increased the synaptic coupling’s strength from inhibitory to excitatory neurons (J_ei_) from −0.75mV to −0.5mv in steps of 0.025 mV. Note that within each simulation, J_ei_ and θ _inh_ were kept constant, but varied across simulations as the EI ratio was systematically changed.

For simplicity, we fix the excitatory firing threshold (θ_exc_) at 20 mV and set the synaptic coupling strengths within the network as follows: excitatory to excitatory neurons (J_ee_) = 0.1 mV, excitatory to inhibitory neurons (J_ie_) = 0.1 mV, inhibitory to inhibitory neurons (J_ii_) = −0.5 mV. J_ei_ varies between −0.75mV to −0.5mV in steps of 0.025 mV.

### Simulation Parameters

Simulations were written in the Python programming language, using the Brian2 simulator (69). The differential equation, as defined in equation (1), was solved using the forward Euler integration method with a time step size of 0.1 ms (21,70). Simulations were run for 1s.

To evaluate the impact of the EI ratio on the network dynamics, we initially measured the network firing rate, variability in the timing between consecutive neural spikes and the synchrony between neurons in the networks as explained in the following section.

### Analysis

#### Network Firing Rate

In order to obtain the overall activity level and functional state of the network, we measured the average population firing rate:

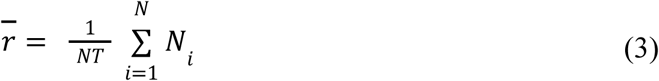

where N_i_ denotes the number of spikes for neuron i, T = 1s and N is the number of neurons whose inter-spike intervals (ISI) were measured.

Due to the high computational cost of analyzing the full neural population, we randomly sampled N = 500 neurons from both excitatory and inhibitory populations, preserving the overall excitatory-inhibitory (EI) ratio. To avoid overlap between simulated and analyzed populations—since direct external input can induce synchronization in excitatory neurons—we selected excitatory neurons that did not receive external stimulation.

#### Coefficient of Variation of the Inter Spike Interval (CV_ISI_)

To measure the variability in the timing of neural spikes, we quantified the ratio of the standard deviation to the mean of the ISI distribution, as indicated in Equation 4, where 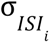 denotes the standard deviation of the ISIs, and 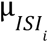 denotes the meaning of the ISIs (71). Here, N denotes the total count of neurons in the network for which the inter-spike intervals (ISI) were measured. we measured the CV _ISI_ of a chosen N= 500 neurons within the network:

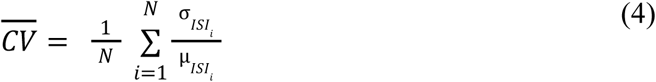

CV _IS_ can take values between 0 and 1. A value of 0 indicates highly regular spike trains, with consistent intervals between spikes. Conversely, a value of 1 signifies highly irregular firing patterns, with large variations in the timing of spikes.

#### Synchronicity within the network

To determine the synchronicity between neurons in the network, we discretized the spike train into 10 ms time-bins and calculated the cross-correlation over the discretized spike trains x(t) of the 500 chosen neurons (as described above) as follows:

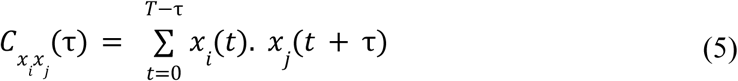

Where 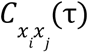 is the cross-correlation between the spike trains of neurons *x_i_* (*t*) and *x_j_*(*t*) at time lag τ.

In order to measure the overall synchrony and connectivity within the network, we need to average the maximum normalized cross-correlations for all pairs of neurons in a network as follows:

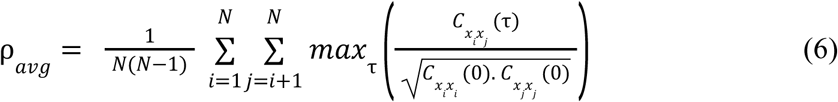

where N is total number of neurons used for cross-correlation (N=500) and 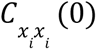 and 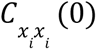 are the auto-correlation functions of the spike trains of neurons *ii* and *jj* at zero lag, which were used to normalize the cross-correlation values 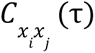 between 0 and 1, in order to make them comparable across different pairs of neurons. 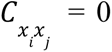, indicating absence of synchrony and 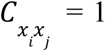 denoting perfect synchrony. For each pair of neurons, we determine the maximum value of the normalized cross-correlation across all time lags τ. This represents the peak synchronization between the two neurons, irrespective of the specific time lag. The average of these peak synchronization values across all neuron pairs is then calculated. This choice of correlation method was motivated by the possibility of lags in the co-activation between neurons and therefore we did not want to focus on a 0-lag measure. In reality, the delay of the peak in the cross-correlation relative to 0-lag was only 3 ms +/- 3 ms. We did not use an integrated measure, because we expect that this would ‘wash out’ the results, if there is an overall high level of correlation, which would overwhelm the peak correlations.

#### Participation Ratio (PR)

To assess the network’s dimensionality, we use the PR (5,40) This metric relies on a straightforward eigenvalue-based formula

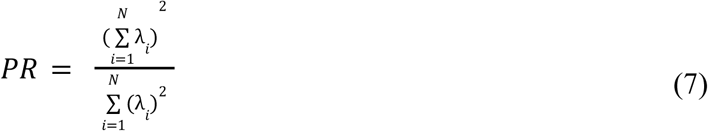

where λ*_i_* are the eigenvalues of the covariance matrix of the firing rates obtained in response to the successive presentation of the two stimuli. We found this to be a good aggregate measure that explains the differences in the function generation capacities of networks in different EI ratios (Fig. 3,5).

If the leading eigenvalue carries all the variance, then PR = 1. At the other extreme, if all eigenvalues are equal, the variance is spread evenly across all the dimensions, and PR = N. The actual value of PR interpolates between these two extreme conditions to estimate the intrinsic dimensionality.

### Stimulus Decoding

We used the Support Vector Machine (SVM) as our supervised decoding model to decode the two time-dependent stimuli from spike trains of the neural network model. The decoder was implemented with a radial basis function kernel SVM, which is particularly suitable for decoding tasks involving high dimensional input features, such as neuronal firing rates. The implementation was done in scikit-learn, a widely-used machine learning library in Python. The performance of the decoder was evaluated using 5-fold cross validation. In this approach, the data was divided into training and test sets. We used 80% of data as the training set, which were selected randomly. The remaining 20% served as the test set to predict the presented stimulus identity in each trial. We analyzed a total of 30 trials per stimulus. The SVM decoder was trained on a training set to distinguish between two time-dependent stimuli. After training, the decoder returned a vector of predicted class labels, based on the trained SVM model. We used the classification accuracy, by comparing the results of the SVM predictions with the actual data to estimate the generalization error. We quantified decoding performance as the percentage of correct predictions on the test set. Considering 2 stimuli, the chance level is 50%.

### Analyzing Electrophysiology Dataset from Visual Cortex

To evaluate the impact of the EI ratio on the dynamics and computational capacity of the spiking neural network simulations, we compared our findings with spiking activities recorded from the mouse visual system. We analyzed a publicly available dataset from the Allen Brain Institute as follows: The dataset from the Allen Brain Observatory was accessed through the AllenSDK (39). The data used in our analysis comprised the Brain Observatory dataset (referred to as dataset 1, 32 sessions). During the recordings, mice were exposed to a range of stimuli, including natural scenes, each with varying durations and presentation sequences across the datasets.

Each recording measured activity in multiple brain regions (e.g., visual cortex, hippocampus, thalamus, midbrain) using up to 6 Neuropixels probes simultaneously (374 data channels, sampling rate 30 kHz) (72). The Allen Brain Institute performed the spike sorting and registration procedures using Kilosort2 (73) and included units that passed the default quality standards. We excluded any invalid intervals marked as not a number (NaN). For more detailed information about experiment preparation, visual stimuli, data acquisition, and data pre-processing steps, please refer to the Technical White Paper from the Allen Brain Observatory on Neuropixels Visual Coding.

In this study, we selected units in the primary visual cortex (VISp) that were stimulated by a specific stimulus, namely "natural scenes," as they contain more visual information and have a higher capacity for decoding images from neural activity (46). We investigated the role of the EI ratio composition across cortical layers. The activity was recorded from the onset of each stimulus block presentation. These presentations consisted of 118 natural images, each repeated 50 times, taken from the Berkeley segmentation dataset (74), the van Hateren natural image dataset (75) and the McGill Calibrated colour image database (76). The images were presented in greyscale and were contrast-normalized and resized to 1,174 × 918 pixels. The images were presented in a random order for 250 ms each. For analysis, we considered neurons with a signal-to-noise ratio of more than 2 and partitioned their spiking activity into bins of 10 ms each.

From the recorded cortical layers, we classified the cell types in layer 4 and layer 2/3 into excitatory and inhibitory based on the distributions of their AP waveform durations. We used a threshold of 0.43 ms (S2 Fig) to classify the neurons. Cells with AP waveform durations shorter than 0.43 ms were classified as inhibitory, while those with durations longer than 0.43 ms were classified as excitatory. We found 241 excitatory neurons and 81 inhibitory neurons in layer 4 of VISp, and 181 excitatory neurons and 54 inhibitory neurons in layer 2/3 of VISp. To compare our analysis with the barrel cortex, we maintained similar EI ratios in layers 4 and 2/3. et al. (2023) (46) reported that layer 4 of the barrel cortex has an EI ratio of 7.3, with a balance of 1374 excitatory neurons and 187 inhibitory neurons per column. Layer 2/3 had an EI ratio of 5.25, with 2232 excitatory and 425 inhibitory neurons per column.

To achieve an EI ratio of 5.3 in layer 2/3, we selected all 181 excitatory neurons and randomly selected 34 inhibitory neurons from the 54 available in layer 2/3. To maintain a similar inhibitory neuron ratio between layers 2/3 and 4 as observed in the barrel cortex (1.62), we randomly selected 111 neurons from the 241 excitatory neurons in layer 4 of VISP. Finally, to preserve an EI ratio of 7.35 in layer 4, we randomly selected 15 inhibitory neurons from the 81 available in layer 4 of VISP. All random selections were repeated three times for further analysis.

To estimate the dimensionality of the selected neurons in layer 2/3 and layer 4 of VISp, we used the PR (Equation 7) on the covariance matrix of the firing rates of neurons observed in response to natural scene stimuli.

### Natural scenes decoding

As described for the model, we used a SVM decoder model to measure how neurons in layer 2/3 and layer 4 encode natural scenes. The implementation of the SVM here is the same as the SVM model in decoding two time-dependent stimuli from spiking neural network models: we used 5-fold cross validation with 80% of the data as training set and remaining 20% as the test sets. The SVM decoder was trained on a training set to distinguish between 118 natural images. After training, the decoder returned a vector of predicted class labels, based on the trained SVM model. We used the classification accuracy, by comparing the results of the SVM predictions with the actual data, to measure the encoding capacity of the layer 2/3 and layer 4 to decode the neural images. Considering 118 natural images, the chance level is 0.0085.

### Data availability statement

All raw and processed data and analysis scripts are stored in the repository of the Donders Institute for Brain, Cognition and Behaviour (https://doi.org/10.34973/nnr8-eh03) and can be obtained from the author upon reasonable request.

Publicly available Allen datasets were analyzed in this study. This data can be found at: https://portal.brain-map.org/circuits-behavior/visual-coding-neuropixels

### Relation to analytical work

Note that our initial model state closely aligns with the configuration of Ostojic et al. (2014) (22). We do not observe state transitions in our work, and hence an analytical analysis into such state transitions would not show new insights. The observed increases or decreases in computational capacity and dimensionality are within the inhibition dominated balanced state. Therefore, we decided not to repeat the mathematical analysis already thoroughly addressed in the Ostojic et al. (22) study and instead focused our efforts on evaluating how the excitatory-inhibitory (EI) ratio influences the network’s coding capacity.

## Acknowledgements

FZ and AA acknowledge funding from NWO Vidi grant VI.Vidi.213. 137. BE acknowledges funding from NWO Vidi grant 016.Vidi.189.052.

## Supporting information

**Fig. S1:**
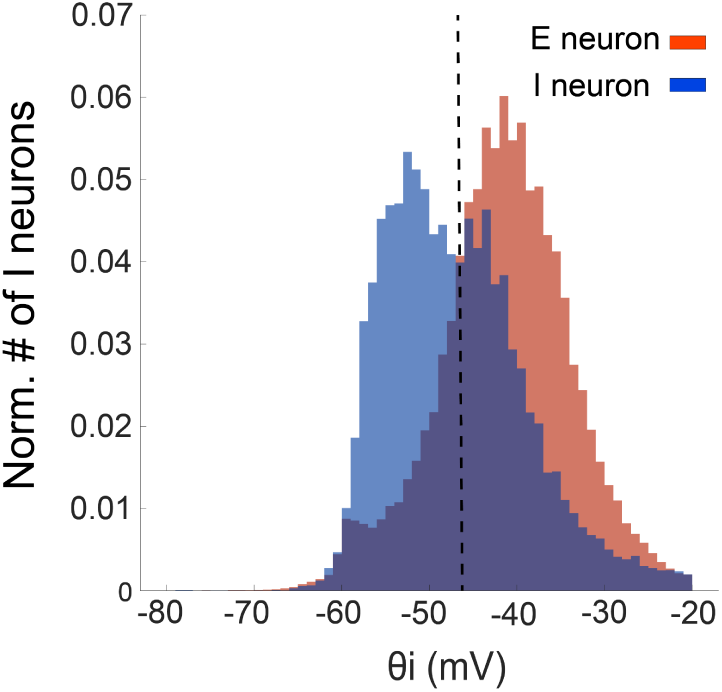
Distribution of firing threshold in inhibitory and excitatory neurons of the mouse barrel cortex. The dashed line indicates the threshold for differentiating between the two cell types, revealing that inhibitory neurons generally have lower firing thresholds than excitatory neurons (data from (38))

**Fig. S2:**
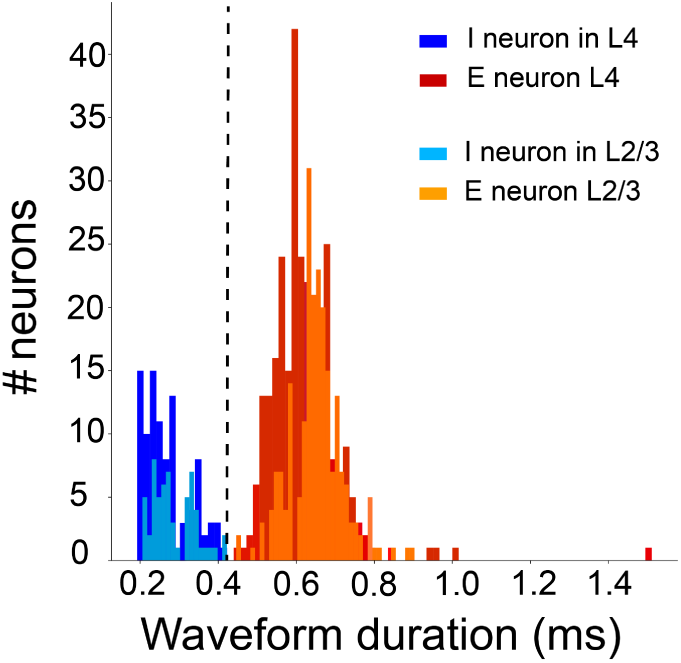
Classification of mouse primary visual cortex neurons based on AP waveform duration. The dashed line represents the threshold for distinguishing excitatory from inhibitory cells (data from (39))

**Fig. S3:**
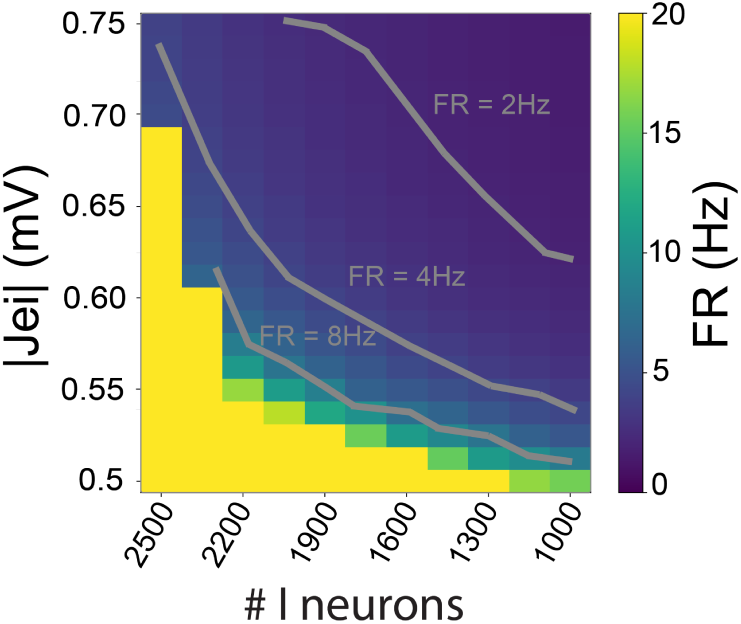
Firing rate of the network across varying the number of inhibitory neurons N_inh_ (between 2500 and 1000), and consequently the EI ratio (between 4 and 10), the synaptic coupling strength J_ei_ (between −0.75 mV and −0.5 mV), the inhibitory firing threshold θ_inh_ = 20 mV. The figure displays three firing rates: a green line at 8 Hz, a brown line at 4 Hz, and a purple line at 2 Hz, which remain consistent across all EI ratios and Jei values due to a constant inhibitory drive.

